# Murine intrarectal instillation of purified recombinant *C. difficile* toxins enables mechanistic studies of pathogenesis

**DOI:** 10.1101/2020.08.30.274738

**Authors:** Nicholas O. Markham, Sarah C. Bloch, John A. Shupe, Erin N. Laubacher, M. Kay Washington, Robert J. Coffey, D. Borden Lacy

## Abstract

*Clostridioides difficile* is linked to nearly 225,000 antibiotic-associated diarrheal infections and almost 13,000 deaths per year in the United States. Pathogenic strains of *C. difficile* produce toxin A (TcdA) and toxin B (TcdB), which can directly kill cells and induce an inflammatory response in the colonic mucosa. Hirota, *et al*. first introduced the intrarectal instillation model of intoxication using TcdA and TcdB purified from VPI 10463 and 630 *C. difficile* strains. Here, we expand this technique by instilling purified, recombinant TcdA and TcdB, which allows for the interrogation of how specifically mutated toxins affect tissue. Mouse colons were processed and stained with hematoxylin and eosin (H&E) for blinded evaluation and scoring by a board-certified gastrointestinal pathologist. The amount of TcdA or TcdB needed to produce damage was lower than previously reported *in vivo* and *ex vivo*. Furthermore, TcdB mutants lacking either endosomal pore-formation or glucosyltransferase activity resemble sham negative controls. Immunofluorescent staining revealed how TcdB initially damages colonic tissue by altering the epithelial architecture closest to the lumen. Tissue sections were also immunostained for markers of acute inflammatory infiltration. These staining patterns were compared with slides from a human *C. difficile* infection (CDI). The intrarectal instillation mouse model with purified recombinant TcdA and/or TcdB provides the flexibility needed to better understand structure/function relationships across different stages of CDI pathogenesis.

## Introduction

*Clostridioides difficile* infection (CDI) is a potentially life-threatening cause of antibiotic-associated diarrhea and is the most common nosocomial disease in the United States (1, 2). The incidence of recurrent CDI, or repeat infection in the same patient, is increasing at an alarming rate and has contributed to the CDC’s designation of CDI as an urgent threat (3, 4). Widespread infection control and antibiotic stewardship programs have helped reduce the burden of hospital-acquired CDI, but community-acquired infections are becoming more common (5, 6). A better fundamental understanding of CDI pathogenesis is needed to formulate new preventive and therapeutic strategies (7).

*C. difficile* is a Gram-positive, spore-forming bacterium with obligate anaerobic metabolism (8). CDI is a toxin-mediated disease primarily driven by *C. difficile* toxin A (TcdA) and toxin B (TcdB) with less potent contribution by the binary transferase toxin (CDT) (9–11). TcdA and TcdB have intracellular glucosyltransferase activity targeting RhoA, Rac1, and CDC42 (12, 13). This inhibitory modification of Rho family GTPases leads to actin depolymerization and disruption of cell-cell junctions (14, 15). Although TcdA and TcdB share 49% amino acid sequence identity and 69% similarity, they have different potencies and mechanisms in multiple animal models (16). When directly compared in porcine explants, both TcdA and TcdB caused glucosyltransferase-dependent apoptosis, but only TcdB could induce a glucosyltransferase-independent cell necrosis at higher concentrations (17–19).

Here, we describe how purified recombinant TcdA and TcdB affect mouse colon after intrarectal instillation. Importantly, our optimized methodology reveals acute inflammatory colitis with low concentrations of either TcdA or TcdB. Whereas these low concentrations induce mild colitis, more aggressive pathology mimicking severe human CDI is observed at higher concentrations of TcdB. The instillation of TcdB mutant toxins reveals the critical roles of endosomal pore formation and glycosylation for initiating *in vivo* mucosal injury in the mouse. Immunofluorescent images reveal how wild-type TcdB disrupts the luminal surface epithelium. Combinations of low-quantity TcdA and TcdB induce acute inflammatory infiltration that is detected with Ly6G immunohistochemistry (IHC) more accurately than H&E.

## MATERIALS AND METHODS

### Animals and housing

The Institutional Animal Care and Use Committee at Vanderbilt University Medical Center approved this study. Our laboratory animal facility is AAALAC-accredited and adheres to guidelines set forth in the Guide for the Care and Use of Laboratory Animals. The animals’ health was assessed daily, and moribund animals were humanely euthanized by CO_2_ asphyxiation followed by cervical dislocation. C57BL/6J mice (all females, age 8-10 weeks) were purchased from Jackson Laboratories (Bar Harbor, ME) and allowed to assimilate for 1 week to the new facilities, thus avoiding stress-associated responses. Mice were kept in a pathogen-free room with clean bedding and free access to food and water. Cage changes were performed bi-weekly. Mice had 12-hour cycles of light and dark.

### Recombinant *C. difficile* toxin purification

TcdA and TcdB were expressed and isolated as previously described (20, 21). Plasmids encoding His-tagged TcdA (pBL282) or TcdB (pBL377) were transformed into *Bacillus megaterium* per manufacturer’s protocol (MoBiTec). Six L of Luria-Bertani media supplemented with 10 mg/L tetracycline were inoculated with an overnight culture to an OD_600_ of ∼0.1. Cells were grown at 37 °C and 220 rpm. Expression was induced with 5 g/L of D-xylose once cells reached an OD_600_ of 0.3-0.5. After four hours, cells were centrifuged and resuspended in 20 mM Tris, pH 8.0, 500 mM NaCl and protease inhibitors. An EmulsiFlex C3 microfluidizer (Avestin) was used at 15,000 psi twice to generate lysates. Lysates were then centrifuged at 40,000 g for 20 min. Supernatant containing toxin was passed through Ni-affinity column (HisTrap FastFlow Crude, GE Healthcare) initially. Further purification was performed with Q-sepharose anion exchange chromatography (GE Healthcare) and gel filtration chromatography in 20 mM HEPES, pH 6.9, 50 mM NaCl.

Expression and purification of mutants were performed as for wild-type TcdB. L1106K (pBL682) and DxD (pBL765) mutant TcdB toxins were produced as previously described (19, 22). Q5 High Fidelity PCR polymerase (M0491) and NEBuilder HiFi DNA Assembly Master Mix (E2621) were used for PCR reactions and cloning, respectively.

### Intrarectal instillation and sample collection

Toxins were prepared in a total volume of 200 μL per instillation. Mice were anesthetized with isofluorane and confirmed to be sedated by toe pinch. The 200 μL instillation was performed over 30 sec with a flexible plastic gavage applicator (20 gauge x 30 mm) while lightly pinching closed the anus, which was held for an additional 30 sec as previously described (23). Mice were recovered in their cages. After 4 hours, mice were humanely euthanized by CO_2_ inhalation. The colon was isolated and dissected away from surrounding visceral tissue. The whole colon was flushed and washed in chilled, sterile 1x PBS before rolling into a Swiss-roll and fixing in 10% Formalin at 4 °C for 16 hrs. Tissue was then washed in 1x PBS and stored in 70% ethanol prior to paraffin embedding by the VUMC Translational Pathology Shared Resource. Samples of human *C. difficile*-infected colon tissue were obtained by the Cooperative Human Tissue Network from consented, de-identified donors under the Vanderbilt Institutional Review Board-approved protocol #031078.

### Statistical analysis

Statistical testing and graphical representations of the data were performed using the following packages: ggplot2, ggpubr, ggsignif, rstatix, and tidyverse in R version 3.6.3.(24–29) For image quantification, at least 10 fields of view at 20x magnification were averaged per animal. Statistical significance was set at a *p* ≤ 0.05 for all analyses (*, *p* ≤ 0.05; **, *p* ≤ 0.01; ***, *p* ≤ 0.001; ****, *p* ≤ 0.0001). The Mann-Whitney-Wilcoxon rank sum (Mann-Whitney) test was used to compare two groups, or the Kruskal-Wallis test was used to calculate significance with *post-hoc* analysis using Dunn’s test when two groups were compared within multiple comparisons.

## RESULTS

### Purified recombinant TcdB reproduced human pathology

To test whether intrarectal instillation of purified recombinant *C. difficile* toxin results in pathology similar to the human disease, we compared colonic mucosa from this mouse model to human CDI. Separate blocks of formalin-fixed, paraffin-embedded tissue were obtained from a patient who died from complications related to severe CDI. H&E staining shows massive acute inflammation, edema, and epithelial injury (Fig. 1A, top row). Four hours after instillation of 50 μg TcdB, the distal mouse colon was removed and prepared for H&E staining, which showed focal acute inflammation, edema, and apoptosis similar to the severity seen in human CDI (Fig. 1A, bottom row). Pseudomembrane formation, a hallmark of CDI pathology, was seen even at this early time point (black arrowheads, Fig. 1A).

**Figure 1:**
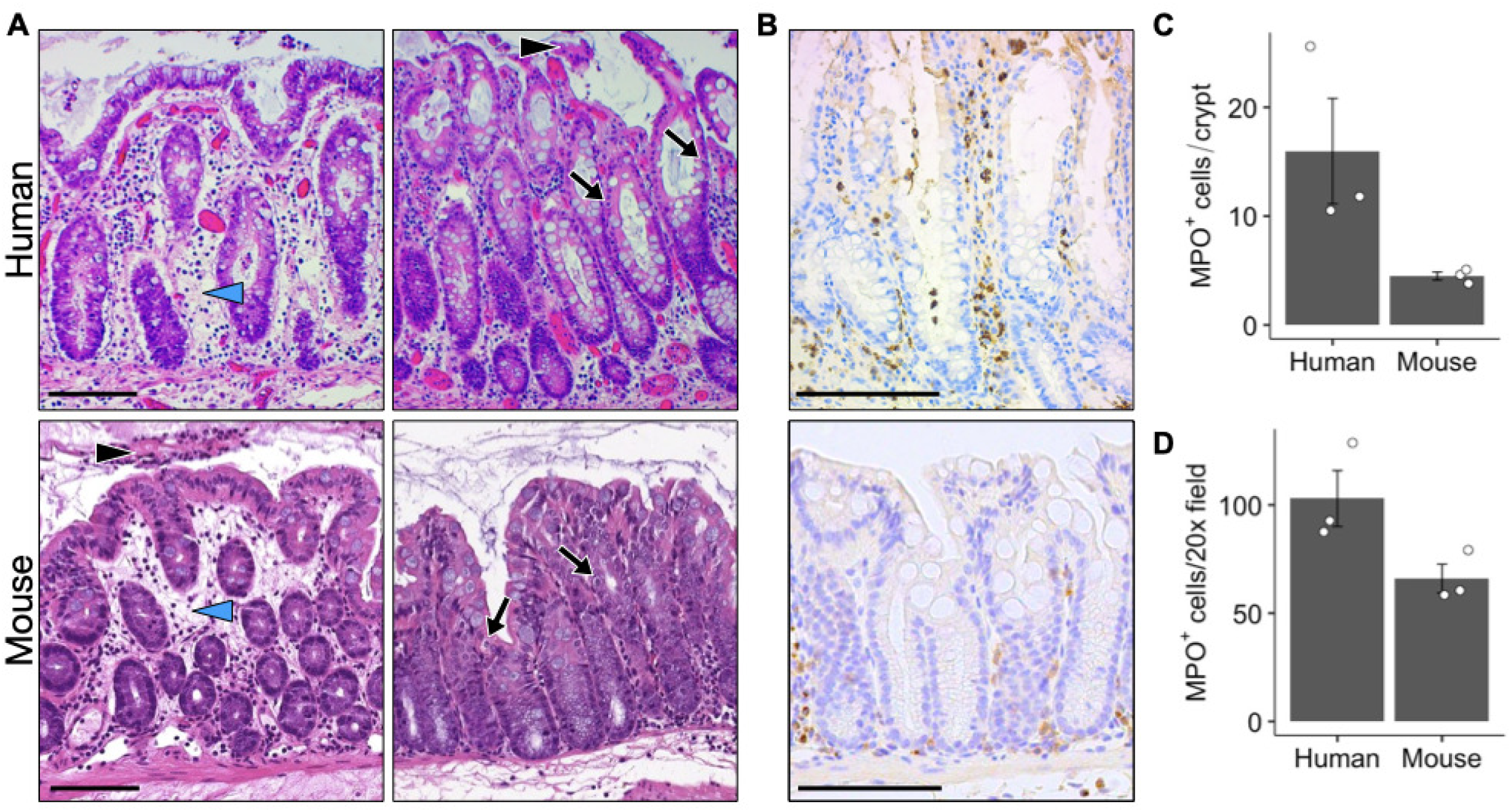
Intrarectal instillation of purified recombinant TcdB recapitulates the histopathology of human CDI. **(A, top row)** H&E images of colon epithelium from a patient with severe CDI shows acute inflammatory infiltration and edema (blue arrowhead), epithelial cell apoptosis (arrows) and pseudomembrane formation (black arrowhead). **(A, bottom row)** Distal mouse colon four hours after instillation of recombinant TcdB (50 μg/200 μL) exhibits acute inflammatory infiltration, edema (blue arrowhead), epithelial cell apoptosis (arrows), and pseudomembrane formation (black arrowhead). **(B)** IHC with anti-myeloperoxidase (MPO) antibody from human CDI colon (top) and mouse colon instillation with 5 μg/200 μL TcdB (bottom). All images within each panel are the same magnification; scale bars = 100 μm. **(C and D)** Quantification of MPO+ cells in human CDI versus mouse colon from panel (B). Error bars show standard error of the mean (SEM); groups were not significantly different by the Mann-Whitney test (*p* = 0.1 for both C and D).

To highlight the neutrophil infiltration for comparison to human CDI, both sets of tissue were stained with IHC using anti-myeloperoxidase (MPO) antibodies (Fig. 1B). In focal areas of mouse tissue, MPO^+^ cells infiltrated the lamina propria and epithelium similarly to the human CDI tissue (Fig. 1B). Across the whole distal colon there appeared to be fewer total MPO^+^ cells per crypt, which may reflect the relatively larger size of human crypts (Fig. 1C). Calculating the number of MPO^+^ cells per 20x field of view appears to better represent the images (Fig. 1D). The histopathological damage and immune infiltration seen here support intrarectal instillation of 50 μg TcdB as a model for human CDI.

### TcdB induced colitis and was concentration-dependent

We chose to focus primarily on TcdB because pre-clinical models of CDI with isogenic knockout strains have shown how TcdB is required for inducing wild-type levels of tissue damage and mortality (10, 11, 30). Furthermore, clinical epidemiological data have revealed a worldwide emergence of *C. difficile* strains with only TcdB and not TcdA or transferase toxin (31–34). Hirota *et al*. first developed the *C. difficile* intrarectal intoxication model using toxins purified from laboratory culture supernatants through dialysis tubing (23). Using recombinant, chromatography-purified TcdB, we performed intrarectal instillations in C57Bl/6J mice with a total volume of 200 μL. Notably, we instilled a higher total volume than the original model (100 μL), because the potential pipetting error or anal leakage of 10% would result in a lower absolute amount of toxin lost. A blinded, board-certified gastrointestinal pathologist scored the edema, inflammation, and epithelial injury according to published criteria (35). In contrast to Hirota *et al*., we saw significant edema and inflammation with 5 μg TcdB (Fig. 2A, B). This quantity also appeared to induce epithelial injury, but this histopathological sub-score did not reach statistical significance compared to vehicle control (*p* = 0.07). Even 0.5 μg TcdB instillation appeared to induce edema and inflammation, but again the histopathological scores were not significantly different from control (*p* = 0.14 and *p* = 0.08, respectively). Increasing the quantity of TcdB induced increased damage in all three sub-scores, with 25 μg and 50 μg TcdB having had the largest effect (Fig. 2A, B). These two treatments resulted in similar histopathological scores, but only mice in the 50 μg TcdB group developed early pseudomembrane formation (12.5%, n = 2/16) composed of sloughed epithelial cells, neutrophils, and fibrin debris in the lumen (Fig. 2A, black arrowhead). With regards to the edema score, it was frequently difficult to differentiate mild edema from tissue sectioning artifact when space was seen between the lamina propria and muscularis mucosa. None of the mice had physiological changes, such as hunched posture or diarrhea, at the 4-hour time point used in this study. In the course of these experiments, we directly compared phosphate buffered saline (PBS) and Hank’s Buffered Salt Solution (HBSS) as diluents for the purified toxins. Unpublished work from our lab suggested there might be a buffer-dependent difference in TcdB potency *in vitro*; however, with 5 μg TcdB, either diluent resulted in the same effect (Supp. Fig. 1A). Similarly, cefoperazone pre-treatment, which is commonly used in the *C. difficile* spore gavage mouse model, did not cause any significant difference in histopathology compared to water-only controls (Supp. Fig. 1B). The edema and inflammation observed with 5 μg TcdB and its dose-dependent effect highlight the robustness of this model for studying mild to severe disease.

**Figure 2:**
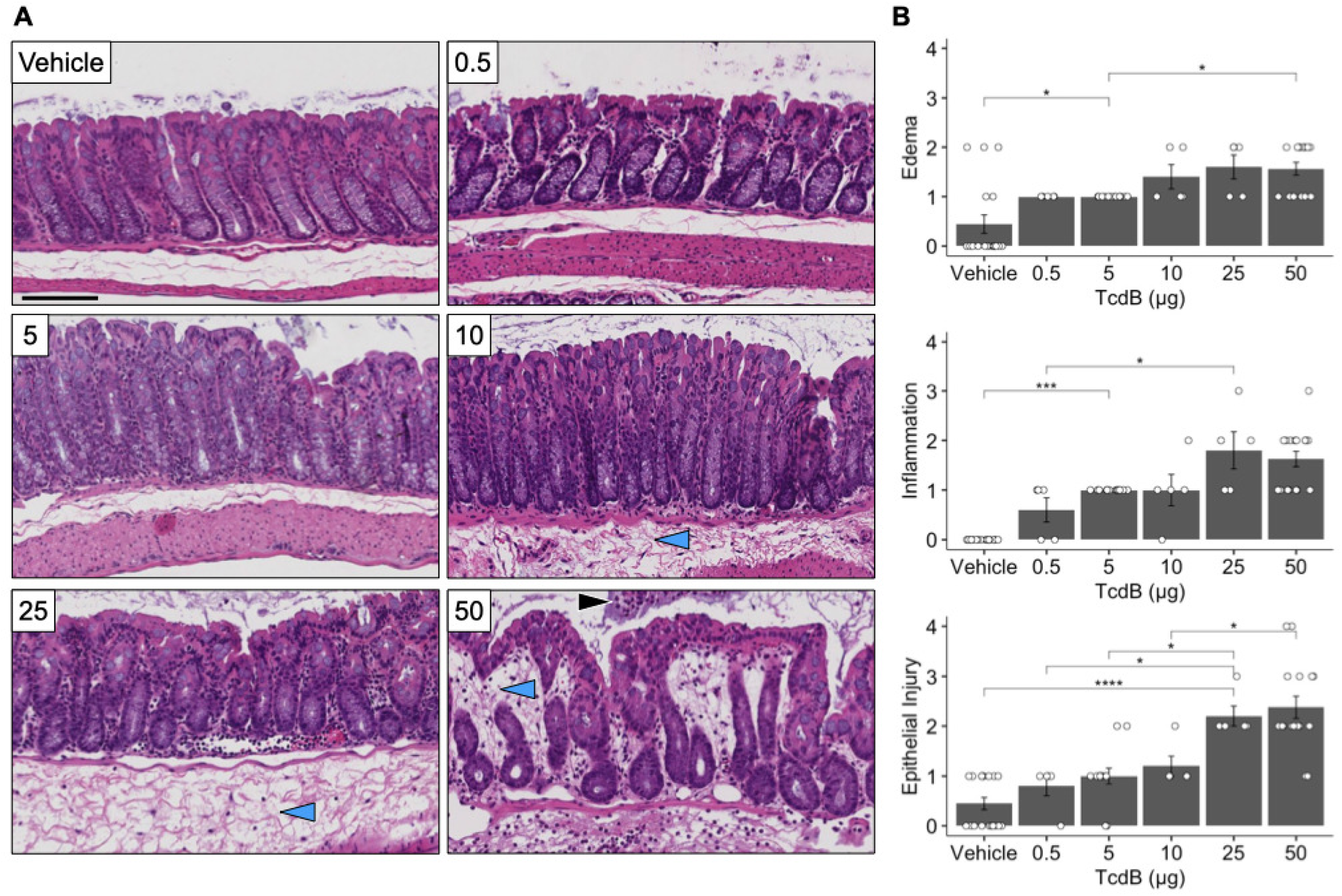
Purified recombinant TcdB induces colitis at low doses and is dose dependent. **(A)** Four hours after intrarectal instillation of TcdB (total µg/200 µL shown as inset), mouse colons were prepared for H&E as in Fig 1. Vehicle negative control was either PBS or HBSS. Representative images are shown from n = 5 - 18 mice per group. Pseudomembrane formation (black arrowhead), edema (blue arrowheads). All images are the same magnification; scale bar = 100 µm. **(B)** Histopathological scoring by an expert gastrointestinal pathologist blinded to the conditions was recorded as mean score per group; error bars = SEM. All three sub-scores were significant by the Kruskal-Wallis test with large effect size, and *post-hoc* analysis with Dunn’s test generated *p* values * ≤ 0.05, ** ≤ 0.01, or *** ≤ 0.001 as indicated. Not all significant comparisons are shown with brackets.

### Mutant TcdB instillation revealed structure/function relationship

To determine the functional impact of specific TcdB components, we performed intrarectal instillation with 50 μg of either wild-type, L1106K, or D286/288N (referred to here as DxD) TcdB. The L1106K mutation ablates a hydrophobic region in the TcdB translocation domain, thus prohibiting pore formation in intracellular endosomes and the subsequent release of the glucosyltransferase domain into the cytosol (22, 36). The DxD motif mutation blocks the TcdB glucosyltransferase catalytic site from binding UDP-glucose and manganese co-substrates, therefore inhibiting the enzymatic activity of the glucosyltransferase domain (17, 37–39). Histopathological scoring of H&E stained tissue showed L1106K and DxD mutant TcdB toxins giving similar damage scores compared to vehicle control (Fig. 3). While not statistically different than vehicle control (*p* =0.43), the DxD mutant did appear to induce mild acute inflammation (Fig. 3 and Supp. Fig. 2). The use of recombinant TcdB allows for more facile experiments to test how specific toxin structures contribute to disease pathology.

**Figure 3:**
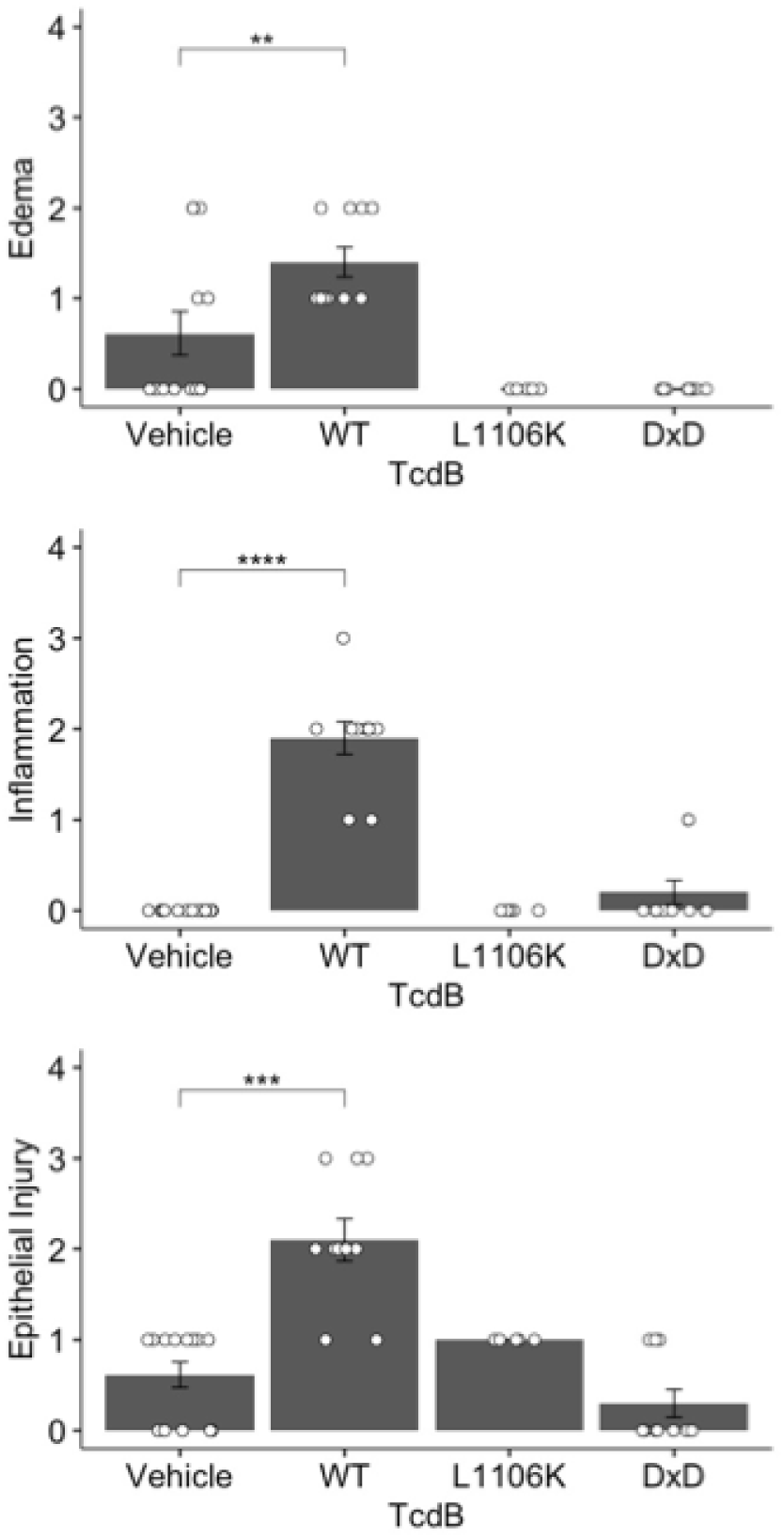
Mutant TcdB toxins lacking pore formation or glucosyltransferase activity fail to induce damage. Recombinant wild-type TcdB from the VPI 10463 reference strain (WT) was produced in *B. megaterium* in parallel with mutant versions L1106K and DxD, which lack endosomal pore formation and glucosyltransferase activity, respectively. Error bars = SEM; n = 5 - 13 mice instilled with 50 μg toxin. All three sub-scores achieved significance by the Kruskal-Wallis test for multiple comparisons with a large effect size, and *post-hoc* analysis with Dunn’s test generated *p* values * ≤ 0.05, ** ≤ 0.01, *** ≤ 0.001, or **** ≤ 0.0001 as indicated. Not all significant comparisons are shown with brackets.

### Histopathological damage occurred near the lumen without crypt base changes

In a CDI mouse model using oral gavage of infectious spores, adherens junction instability was described throughout the crypt axis from base to lumen (40). We used immunofluorescent staining to examine adherens junctions and the apical brush border after intrarectal intoxication. In Fig. 4A, E-cadherin and Villin immunofluorescence were used to highlight TcdB-induced alterations in the colonic epithelial luminal border. Specifically, the luminal invaginations appeared more numerous and larger in size, and the staining intensity within individual cell membranes appeared to vary. Qualitative changes in the amount of membrane protein sodium-potassium ATPase (Na^+^/K^+^-ATP) and sodium-potassium-chloride channel (NKCC) were also observed (Fig. 4B). However, the mean pixel intensity throughout the distal colonic epithelium was unchanged across mouse groups (Fig. 4C). In addition to comparing TcdB from the VPI 10463 reference strain, we also performed instillations with the L1106K mutant. As we have shown in Figure 3, the L1106K mutant has the same histopathological and epithelial cell membrane effect as vehicle negative control (Fig. 4C). These data illustrate how the more luminally located epithelial cells are first affected in the colonic epithelium.

**Figure 4:**
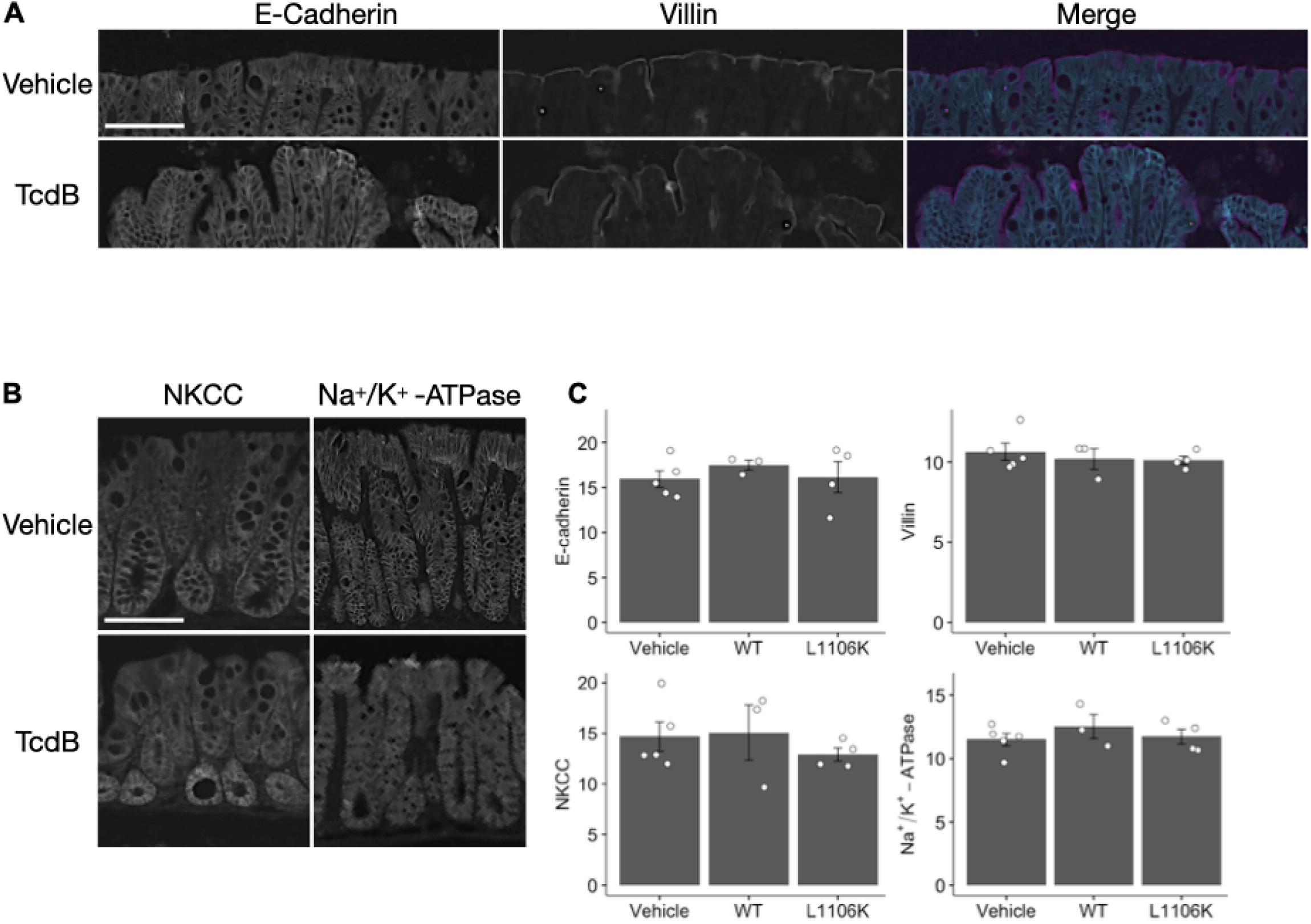
TcdB intoxication affects epithelial architecture but not staining intensity of epithelial cell markers at 4 hours. **(A)** Representative immunofluorescence images of mouse colon showing altered epithelial architecture with more luminal irregularities and invaginations than the vehicle control. TcdB mice were instilled with 50 μg of wild-type TcdB from the VPI 10463 reference strain (WT), and vehicle controls were instilled with PBS only; n = 5 mice/group. **(B)** Representative immunofluorescence images showing no changes in crypt base structure as shown by sodium-potassium-chloride channel (NKCC) and sodium-potassium ATPase (Na^+^/K^+^-ATPase). All images in (A) and (B) are the same magnification; scale bars = 100 μm. **(C)** Immunofluorescence quantification of mean pixel intensity in distal colon epithelium. Error bars = SEM; n = 3 - 5 mice per group. No comparisons were statistically different by the Kruskal-Wallis test for *p* ≤ 0.05.

### TcdA and TcdB combination induced subtle acute inflammation

To continue leveraging our ability to see tissue damage with low quantities of toxin, we sought to describe the effect of combining TcdA and TcdB in this model. Recombinant wild-type TcdA from the VPI 10463 reference strain gene was prepared and purified similarly to TcdB. Both were instilled either separately or together for 4 hours, and tissues were processed by H&E staining and IHC for the Ly6G mouse antigen to identify myeloid-derived cells (41) (Fig. 5A, Supp. Fig. 3). The IHC staining shows more Ly6G^+^ cell infiltration after TcdB alone than with an equivalent quantity of only TcdA (Fig. 5A, B). Moreover, 5 μg TcdB added to either 1 μg or 5 μg TcdA increased the number of Ly6G^+^ cells per crypt (Fig. 5B). These findings suggest TcdB has a greater impact than TcdA on recruitment of neutrophils and macrophages in the mouse colon. We also observed how the histopathological scoring for edema, inflammation, and epithelial injury did not clearly differentiate the groups as well as measuring Ly6G^+^ cells/crypt (Fig. 5A-C).

**Figure 5:**
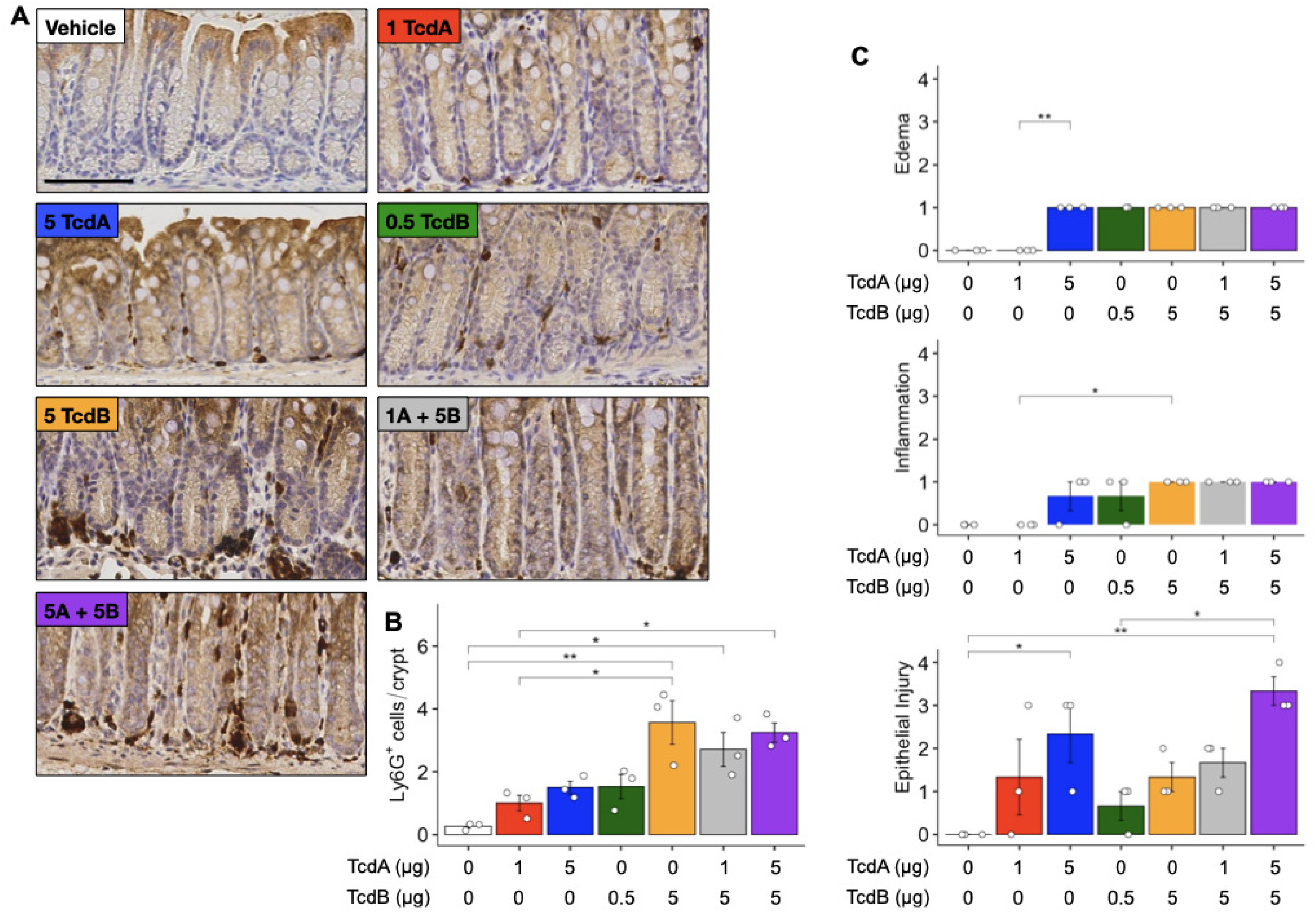
Combination of purified recombinant TcdA and TcdB induces subtle colitis. **(A)** Representative images of IHC Ly6G-stained distal mouse colon. Inset labels indicate μg TcdA and/or μg TcdB intrarectally instilled in the same total volume of 200 μL. All images are the same magnification; scale bar = 100 μm. **(B)** Ly6G^+^ cells/crypt are shown as the mean +/- SEM (n = 3 per group). An average of 13.7 crypts were counted per 20x field of view, and 10 fields were counted per mouse. Kruskal-Wallis test was significant with *p* = 0.0081 and *η*^2^ = 0.811 indicating a large effect size. **(C)** Histopathological scoring as in figures 2 and 3. Brackets indicate * for *p* ≤ 0.05 and ** for *p* ≤ 0.01 by *post-hoc* analysis with Dunn’s test.

## Discussion

Here, we present results from an *in vivo* model for *C. difficile* toxin exposure using purified, recombinant TcdA and TcdB. One benefit of using *in vivo* models compared to tissue explants or organoid cultures is the ability to measure neutrophil infiltration during a fully intact acute inflammatory response. Colonic infiltration of neutrophils is a hallmark of CDI, but the effective role of neutrophils is unclear with regards to their beneficial bacterial containment versus their detrimental escalation of cytokine release (42). We observed similar MPO^+^ cell infiltration between human CDI and mouse intrarectal instillation (Fig. 1B-D). Adding 5 μg TcdA to 5 μg TcdB together did not significantly increase the number of MPO^+^ cells/crypt or cells/field in the mouse (data not shown), nor did this combination increase the Ly6G^+^ cells/crypt compared with 5 μg TcdB alone (Fig. 5B). Ly6G is a marker of murine myeloid derived cells and has been used to measure neutrophil recruitment in CDI models (41, 43, 44). It is a sensitive marker for the acute colitis observed with low quantities of TcdA and/or TcdB and is able to differentiate groups better than histopathological scoring (Fig. 5). These observations suggest TcdB contributes more to acute inflammatory infiltration than TcdA.

We wanted to know the minimal quantity of TcdB needed for inducing colonic pathology in the model, because we have shown TcdB has different, concentration-dependent mechanisms of cell killing (17–19). In previous reports, intrarectal instillation and mouse colonic explants required at least 5 μg TcdA or 10 μg TcdB to induce tissue damage (23, 45). Here, we showed 5 μg purified recombinant TcdA has the same effect as these prior studies, but we saw significant edema and inflammation with only 5 μg TcdB (Figs. 2 and 5). The ability of TcdB alone to produce a range of histopathological severity from subtle neutrophil infiltration to pseudomembrane formation in a 4-hour time period is valuable for studying CDI pathogenesis and testing therapeutic or preventive interventions.

The most significant aspect of this recombinant toxin approach is the ability to interrogate easily how changes in toxin structure affect its function. As an example, we have previously shown how a TcdB mutation (^1595^VNFLQS ->^1596^GFE) abolishing Frizzled binding induces the same histopathological response as wild-type (40). Others have used purified recombinant toxin via systemic injection or on explants to show the L543A mutation renders TcdB unable to undergo cysteine protease cleavage while retaining the capacity to induce histological damage (45, 46). In this report, we more thoroughly characterize intrarectal instillation and further illustrate its capabilities using the L1106K and DxD mutant TcdB toxins (Fig. 3 and 4). Neither of these mutant forms were able to induce significantly greater damage than the vehicle control. The L1106K TcdB has been shown concordantly to lack cell rounding or cell killing effects *in vitro* (22, 36). On the other hand, the DxD TcdB has been used *in vitro* and *ex vivo* to demonstrate the glucosyltransferase-independent mechanism for TcdB-induced necrotic cell death (17–19). It is unclear how 50 μg of DxD TcdB failed to produce significant epithelial injury in the present model, but reasons could include, toxin concentration in colonic tissue being below the threshold for the necrotic effect, and/or species-specific differences affecting the host epithelial response. For example, published comparisons of human and murine colonic explants exposed to TcdB show mouse tissue having decreased susceptibility to tissue damage (45). We previously have found porcine explants have a similar sensitivity to TcdB as human (17).

*C. difficile* toxins have been shown to directly affect cell-cell junctions and intestinal permeability (14, 47–49). Other gastrointestinal pathogens, such as *Bacillus fragilis, Escherichia coli, Klebsiella pneumoniae*, and *Salmonella typhimurium*, disrupt junctional proteins (50, 51). Using the VPI TcdB and L1106K mutant (both 50 μg) for intrarectal instillation, we evaluated the epithelial injury by immunofluorescence. The increased luminal invaginations seen by E-cadherin and Villin immunofluorescence illustrate how wild-type TcdB disrupts the epithelial architecture without broadly affecting the intercellular abundance and localization of apical and basolateral proteins NKCC and Na^+^/K^+^-ATP (Fig. 5A-B). This effect is similar to colonic damage induced by diarrheagenic *E. coli* colonization in mice (51). The E-cadherin staining representing adherens junctions and Villin representing the brush border are largely intact (Fig. 5C). Previously, we showed E-cadherin and β-catenin mislocalization in the spore gavage mouse model occurs after 24 hours (40). Surprisingly, pseudomembrane formation can occur 4 hours post-instillation without dramatically affecting adherens junctions in more basally located cells. This observation suggests top-layer cells closest to the lumen are sloughed from the mucosa initially without significant disruption of the underlying cells, which has implications for how host defense mechanisms function during CDI. Future studies will determine these mechanisms by comparing intrarectal instillations to the *C. difficile* spore gavage mouse model of CDI.

## Acknowledgments

Work in the Lacy lab is supported by **NIH AI095755** and **VA BX002943**. Portions of this work were supported by **R35 CA197570** and **GI SPORE P50236733** to Robert J. Coffey. NIH Training Grant in Gastroenterology **T32 DK007673** and a Harrison Society Research Supplement from the VUMC Department of Medicine funded Nicholas O. Markham. Core Services performed through the VUMC Digestive Disease Research Center supported by NIH grant **P30DK058404**. These results are solely the responsibility of the authors and do not necessarily represent official views of the NIH.

## Author contributions

NOM, SCB, JAS, and DBL conceived of the experiments and planned the design. NOM, SCB, JAS, and ENL carried out the experiments. NOM, SCB, JAS, MKW, RJC, and DBL analyzed the data. NOM wrote the manuscript. NOM, SCB, JAS, RJC, and DBL edited the manuscript.

## Abbreviations

CDI: *Clostridioides difficile* infection
DxD: D286/288N mutant lacking glucosyltransferase activity
H&E: hematoxylin and eosin
HBSS: Hank’s Balanced Salt Solution
IHC: immunohistochemistry
L1106K: mutant lacking pore-forming function
Ly6G: lymphocyte antigen 6G (murine myeloid cell marker)
MPO: myeloperoxidase
Na^+^/K^+^-ATP: sodium-potassium ATPase
NKCC: sodium-potassium-chloride channel
PBS: phosphate buffered saline
SEM: standard error of the mean
TcdA: *C. difficile* toxin A
TcdB: *C. difficile* toxin B
WT: wild-type
VPI: VPI 10463 reference strain (ATCC 43255)

## Figure Legends

**Supp. Fig. 1**: Different diluent buffers or antibiotic pre-treatment show no effect. **(A)** Histopathological scores of mouse colonic tissue 4 hours after TcdB (5 μg) was diluted in either HBSS or PBS and intrarectally instilled; n = 3-5 mice/group. **(B)** Histopathological scores comparing mice pre-treated with either cefoperazone (0.5 mg/ml) or water control for 5 days prior to intrarectal instillation with TcdB (5 μg) in HBSS for 4 hours; n = 5 mice/group. Error bars = SEM. Mann-Whitney testing showed no significant differences between conditions.

**Supp. Fig. 2**: Representative H&E images from Figure 3 using purified recombinant wild-type TcdB and mutants for 4-hour intrarectal instillations. All images are the same magnification; scale bar = 100 μm.

**Supp. Fig. 3**: Representative H&E images from Figure 5 using combinations of purified recombinant TcdA and TcdB for 4-hour intrarectal instillations. All images are the same magnification; scale bar = 100 μm. Inset label shows μg or TcdA and/or TcdB in the same total volume of 200 μL.

